# Action-specific feature processing in the human visual cortex

**DOI:** 10.1101/420760

**Authors:** Simona Monaco, Ying Chen, Nicholas Menghi, J Douglas Crawford

**Affiliations:** Center for Mind/Brain Sciences, University of Trento, Trento, Italy; Centre for Neuroscience Studies, Queens University, Kingston, Ontario, Canada; Center for Vision Research, York University, Toronto, Ontario, Canada M3J 1P3

**Keywords:** Action intention, fMRI, Psychophysiological Interactions, Reentrant feedback, Visual cortex

## Abstract

Sensorimotor integration involves feedforward and reentrant processing of sensory input. Grasp-related motor activity precedes and is thought to influence visual object processing. Yet, while the importance of reentrant feedback is well established in perception, the top-down modulations for action and the neural circuits involved in this process have received less attention. Do action-specific intentions influence the processing of visual information in the human cortex? Using a cue-separation fMRI paradigm, we found that action-specific instruction (manual alignment vs. grasp) influences the cortical processing of object orientation several seconds after the object had been viewed. This influence occurred as early as in the primary visual cortex and extended to ventral and dorsal visual stream areas. Importantly, this modulation was unrelated to non-specific action planning. Further, the primary visual cortex showed stronger functional connectivity with frontal-parietal areas and the inferior temporal cortex during the delay following orientation processing for align than grasping movements, strengthening the idea of reentrant feedback from dorsal visual stream areas involved in action. To our knowledge, this is the first demonstration that intended manual actions have such an early, pervasive, and differential influence on the cortical processing of vision.

## Introduction

Most neuroimaging studies of visual-motor integration have focused on the question of how vision is used to plan movements in a feedforward fashion (Fabbri et al., 2016; Verhagen et al., 2012; for reviews see: Gallivan and Culham, 2015; Vesia and Crawford, 2012). Despite the importance of feedforward processes, feedback is required to combine low-level features, such as line orientation, with high-level concepts, such as affordances. Indeed, the intended action itself often determines which visual details are relevant for the feedforward transformation (Craighero et al. 1999). Action-dependent sensory gating is thought to be controlled by reentrant feedback. For example, reentrant pathways from the oculomotor system to the visual system might explain attentional enhancements near the saccade goal (Moore and Fallah 2001, 2003; Moore and Armstrong 2003). Likewise, reentrant pathways could explain reactivation of the visual cortex during the planning and execution of reaching and grasping movements in the dark (Singhal et al. 2013; Chen et al. 2014; Cappadocia et al. 2016). Reentrant feedback for limb control is potentially complex because the hand-arm system is capable of many types of action and interacts physically with the surrounding world (Perry and Fallah 2017). For instance, intrinsic object features are important for grasping, whereas pointing only requires knowledge of the object location (Van Elk et al. 2010). Further, the same object feature, such as orientation, might be processed differently depending on the nature of the action (Gutteling et al. 2011), e.g., grasping an object versus matching the orientation of an object by aligning the hand. At present, the neural mechanisms for reentrant feedback during action planning, as well as the extent to which the activity in the early visual cortex (EVC) is differentially modulated by specific action plans, object features or both, remain unknown.

The influence of action on perception is well established. For example, placing the hand near a stimulus can influence blindsight (Schendel and Robertson 2004; Brown et al. 2008), extinction (di Pellegrino and Frassinetti 2000), reaction times (Reed et al. 2006), and visual search (Abrams et al. 2008), enhance visual memory recall (Heuer, Crawford, et al. 2016) and enhance orientation selectivity in macaque’s V2 neurons (Perry et al. 2015). Further, neurons in macaque’s anterior intraparietal sulcus (AIP), an area known to be involved in grasping, show enhanced responses to action instruction when a stimulus is presented before as opposed to after the action cue (Baumann et al. 2009). Recently, Gutteling and collegues (2011) have shown that such motor enhancements can be specific to action-relevant features. For instance, orientation perception is more accurate when one intends to grasp an object as opposed to simply point towards it. Importantly, this effect is attenuated by stimulation of the anterior intraparietal sulcus (aIPS), known to have a role in the control of grasp also in humans (Culham et al. 2003; Gallivan, Mclean, Valyear, et al. 2011; Gutteling et al. 2013), suggesting that the aIPS is involved in the reentrant filtering of visual orientation. Some of this reentrant filtering might occur at the level of occipital cortex. For example, lateral occipital cortex (LOC) is more active during delayed grasping than reaching movements in the dark (Singhal et al. 2013), presumably because of the additional sensory processing required to shape the hand for grasping an object. However, little more is known about how the details of motor planning influence visual feature processing, how early this occurs in the visual system, and which reentrant feedback pathways are involved.

It might not be surprising to find action-specific stimulus processing in dorsal visual stream areas because of its well-known association with action planning and execution (Goodale and Milner 1992; Culham et al. 2003). However, based on recent evidence that several occipital areas are involved in action, even in absence of online visual information (Singhal et al. 2013; Chen et al. 2014; Cappadocia et al. 2016), we hypothesized that action-specific plans might influence visual feature processing as early as the primary visual cortex, and therefore might even influence ventral visual stream areas typically associated with object perception. We further hypothesized that if this influence is mediated by recurrent motor feedback, it should result in patterns of task-related functional connectivity between the EVC and hand motor areas. In addition, according to the theory of reverberating circuits (Hebb 1949), the connections between areas in the dorsal visual stream and early visual cortex would persist well after the visual stimulus (the origin for feedforward visual signals) has disappeared.

Here, we tested these hypotheses with functional Magnetic Resonance Imaging (fMRI) by using a cue-separation task in which participants performed one of two actions (manual alignment and grasp) towards a rod in two possible orientations. While the location and orientation of the rod were the same for the two actions, the same orientation required different kinematics for manual alignment vs. grasping movements. To examine the influence of specific action instructions on the cortical processing of vision, we temporally separated action cues and stimulus presentation, with the action being specified *before* the stimulus. Specifically, we instructed subjects to either grasp or align their hand with the oriented object, then showed them the orientation of the object, and finally provided a ‘go’ signal that cued participants to perform the action. We examined if the cortical response following the visual stimulus (both overall activation and orientation specificity) was modulated by the action plan, and used Psychophysiological Interaction (PPI) analysis to identify potential neural pathways for action-related modulation of sensory information.

In sum, while several studies have shown the existence and importance of reentrant (top-down) processing in perception (Reicher 1969; Weisstein and Harris 1974; McClelland and Rumelhart 1981; Coltheart 2011), much less is known about the role of reentrant feedback in action and whether the descending signals reach V1 and modify its activity. Our results suggest that during action planning, reentrant signals from higher cortical areas not only reach V1 but also modulate its activity in a task-dependent manner.

## Material and Methods

### Overview of Experiment and Hypotheses

Fourteen human participants lay on the bed of the MRI and used their right (dominant) hand to perform delayed grasp or hand-alignment actions towards a real 3D rod oriented vertically or horizontally and placed on an apparatus located above their pelvis (**Figure 1A and B**). Our experimental paradigm (**Figure 2A**) exploited a cue-separation task based on previously published neurophysiological (Baumann et al., 2009) and neuroimaging studies (Beurze et al. 2009; Cappadocia et al. 2016). In each trial, participants received an auditory cue (Auditory Action-cue) followed by an 8-s delay (Delay 1) and then viewed the oriented rod (Visual Orientation-cue) followed by another 8-s delay (Delay 2). Participants maintained gaze fixation throughout the trial in an otherwise completely dark space. At the end of the Delay 2, an auditory “go” (Execution-cue) instructed the participants to perform the movement toward the rod. We used a slow event related design to allow the hemodynamic response to return to baseline during the ITI (**Figure 2B**), and randomly interleaved task (Grasp, Align) and stimulus orientation (Horizontal, Vertical) across trials. The Align condition required participants to adjust the hand and wrist posture according to the orientation of the rod, while in the Grasp condition participants adjusted the fingers on the rod in a whole-hand grasp. Therefore, the Align movement required a higher degree of adjustment as compared to the Grasp movement (**Figure 1B**). This paradigm was designed to test whether the task instruction (Align vs. Grasp) modulated the subsequent activation elicited by the visual presentation of the stimulus during Delay 2.

**Figure 1.**
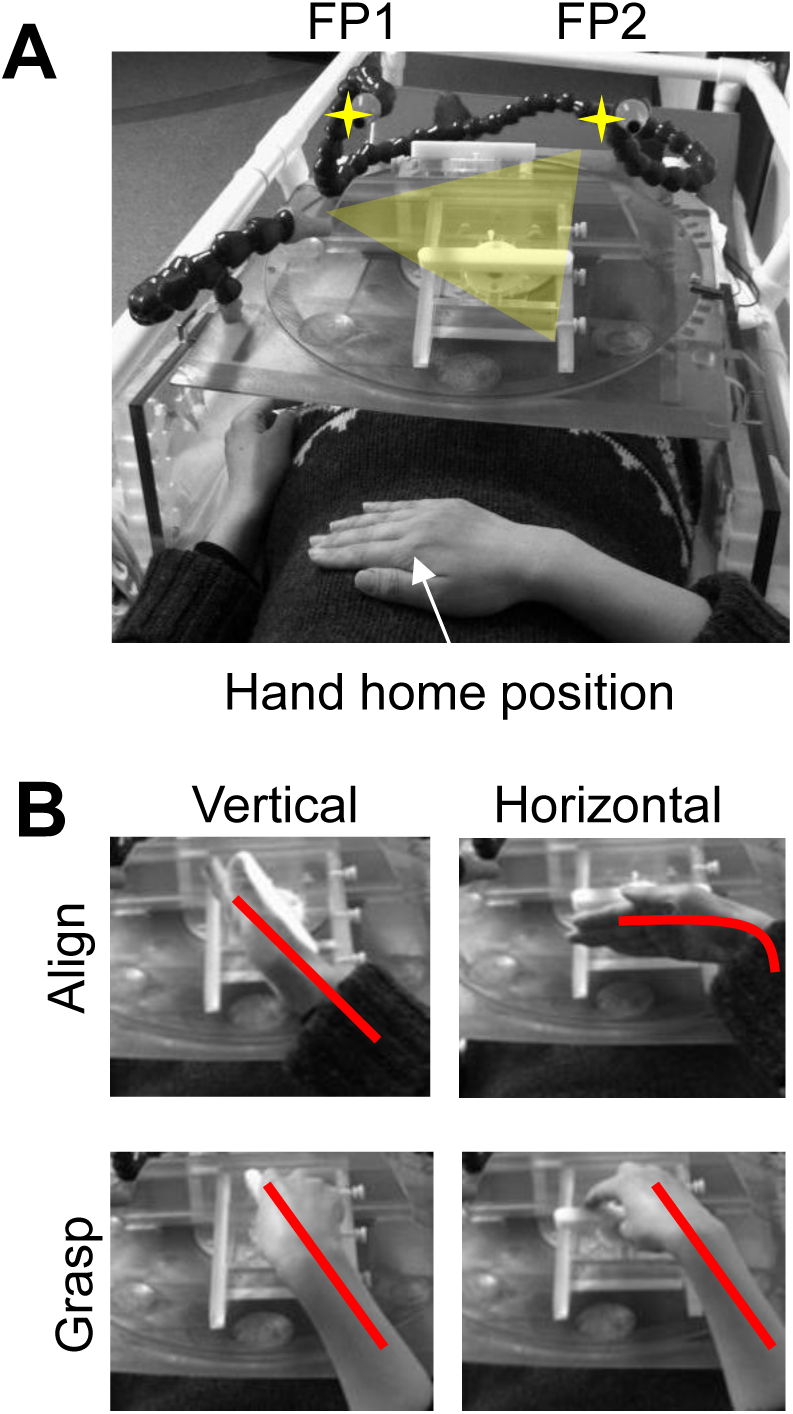
Illustration of the task and setup. Participant performed the tasks (Grasp and Align) towards a rod oriented vertically or horizontally. They were required to gaze at one of two fixation points (FP 1 and 2, marked with a star) for the duration of each trial.

**Figure 2.**
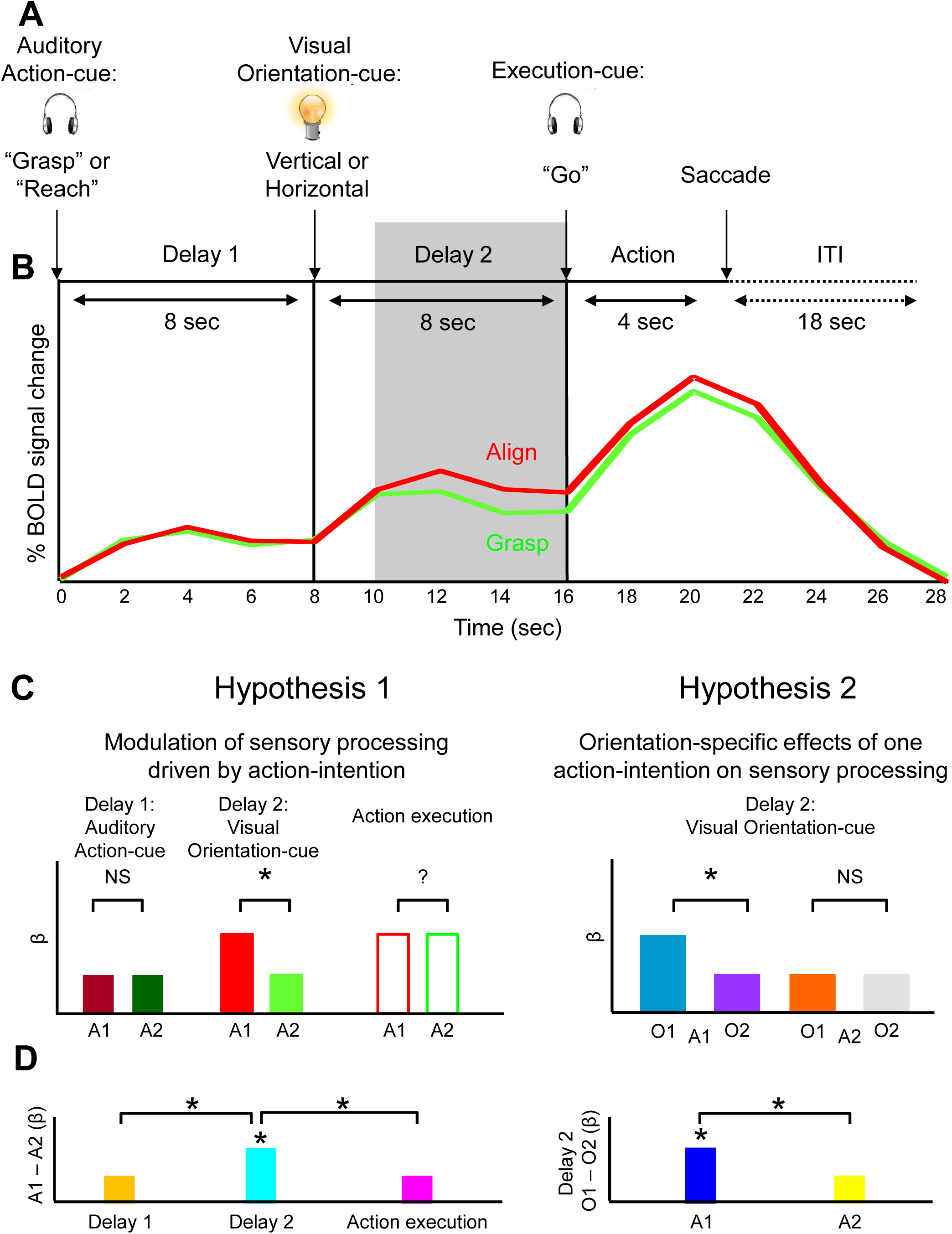
Illustration of the timing and hypotheses. **A.**Each trial started with the auditory cue (“Grasp” or “Reach”) about the action that had to be performed at the end of the trial. After an 8-s delay (Delay 1), the rod was illuminated for 250 ms in one of two possible orientations (Vertical or Horizontal). After another 8-s delay (Delay 2), a “go” cue instructed participants to perform the action towards the rod. After 4 s the fixation cue was turned off for 2 s then re-appeared in the same or different location. We used an inter-trial interval (ITI) of 18 s. **B.** Example of event-related BOLD activity from area PMd in the left hemisphere for Align (in red) and Grasp (in green) conditions. The timecourse shows the average of 14 participants. Events in B are aligned to the corresponding events in A. **C.** Predicted changes in BOLD signal based on our Hypotheses. We hypothesized that the influence of action intention on sensory processing would be reflected in higher activation for Action 1 vs. Action 2 during Delay 2 (upper left panel). In addition, we hypothesized that orientation-selectivity effects would be shown by higher activation for Orientation 1 vs. Orientation 2 in one of the two actions (upper right panel). **D.** Summary of our hypothesis in terms of differences between Action 1 vs. Action 2 in Delay 1, Delay 2 and Action execution (lower left panel) as well as Orientation 1 vs. Orientation 2 in Delay 2 (right lower panel).

**Figure 2C and D** illustrates the general theory and motivation for our analysis using hypothetical β values. Here, our two action instructions are arbitrarily labeled Action 1 and Action 2. Row C illustrates predicted β weights for each of the experimental conditions during Delay 1, Delay 2, and the Execution phase of our task. To compare the differences between conditions, in **Figure 2D** we show the predicted differences between the β weights in row C during the same epochs.

We designed our experiment to test two hypotheses. First, we hypothesized that cortical regions that receive both visual input and reentrant action-specific input would show differences in visual processing of the stimulus for one action as compared to the other (Hypothesis 1**, Figure 2C**, left panel). This would be reflected in higher differential activation for the two action plans in Delay 2 than Delay 1 and Action execution (**Figure 2D**, left panel). Second, we hypothesized that cortical areas involved in gating specific visual input for planning (as opposed to general sensitivity to the action instruction or motor control of the action) would show different feature specificity during the preparation of these actions. This would be reflected in higher activation for one orientation as compared to the other for one of the actions but not the other (Hypothesis 2, **Figure 2C**, right panel). In practice, we expected the Align task to produce higher activation and orientation sensitivity than the Grasp task, because manual alignment involves an explicit orientation instruction, and implies a higher demand on orienting accuracy, while successful grip occurs somewhat automatically and does not require perfect orientation of the hand, as the position of the fingers on the oriented rod can be adjusted after touching the object.

### Participants

Fourteen participants (3 males and 11 females, age range: 24–42 years, average 32 years) participated in this study and were financially compensated for their time. All participants were right-handed and had normal or corrected-to-normal visual acuity. They gave their consent prior to the experiment. This study was approved by the York University Human Participants Review Subcommittee.

### Apparatus and Stimuli

Stimuli were presented to the participants on the inclined surface of a platform placed above the participant’s pelvis. For details about the set up see Monaco et al., 2014. Each participant lay supine in the scanner with the head tilted allowing a direct view of the stimulus. Participants wore headphones to hear auditory instructions about the task that they were to perform at the end of the trial.

The platform was reached by the participant (from inside the bore) to perform the task and by the experimenter (from outside the bore) to change the stimulus orientation between trials. The platform was made of Plexiglas and was fixed to the bore through hooked feet. The location of the platform could be adjusted to ensure that both the participant and the experimenter could reach it comfortably. The head of the participant was tilted by 20° to allow comfortable viewing of the stimuli. The inclination of the platform could also be adjusted to improve the view and the reachability of the object for each participant. A rectangular surface was angled atop the turntable to improve the visibility of the object and its location could be adjusted for each participant (**Figure 2**). A fitted rotating wheel (radius= ~ 4 cm) was embedded on the surface of the platform. The rod was secured to the rotating wheel through fitted pins. Lateral stoppers on the platform served as markers for the experimenter to adjust the orientation of the rod in the dark for the upcoming trial. The orientation of the rod was adjusted to ensure that each participant could reach it comfortably avoiding awkward hand and wrist postures. A cloth was mounted on the ceiling of the magnet bore to occlude the participant’s view of the experimenter.

The stimulus consisted of a rod (10 cm x 1cm x 1 cm) made from Plexiglass and painted white to increase the contrast with the workspace **(Figure 1A**). The angular size of the rod was approximately 8° and was presented at approximately 8-10° of eccentricity in the lower visual periphery. Participants practiced the task and familiarized with the objects for about 5 min prior the experiment.

Except for the brief illuminated presentation of the object, subjects were in near-complete darkness throughout the duration of the trial (only a small dim light was provided by fixation). During the intertrial interval (ITI), and after the participant had performed the task, the orientation of the rod was quickly changed by the experimenter or left in the same orientation for the next trial.

Optic fibers were used to provide a fixation point, to illuminate the workspace, and to cue the experimenter regarding stimuli on upcoming trials. The participant maintained the fixation on one of the two points of light positioned above the object, so that all objects were presented in the left or right participant’s lower visual field. A bright light (illuminator) was used to briefly illuminate the object at the onset of each event of a trial. The illuminator was placed above the participant’s head and shone light onto the object. Another source of light was based at the end of the platform, visible to the experimenter, but not to the participant, to instruct the experimenter about the orientation of the rod on upcoming trials. All of the lights and audios were controlled by a program in MatLab® 7.1 (The MathWorks, Inc., Natick, MA, USA) on a laptop PC that received a signal from the MRI scanner at the start of each trial. The window in the scanner room was blocked and the room lights remained off such that, with the exception of the dim fixation point which remained on for the duration of a trial, nothing else in the workspace was visible to the participant when the illuminator was off. An infrared camera (MRC Systems GmbH) recorded the performance of each participant for offline investigation of the errors, which were excluded from further analysis. The errors were defined as mistakes in the performance of the participants during the task, such as initiating a movement during the delays, performing a grasp in an align condition or vice versa. Less than 15% of total trials were discarded from the analyses due to participants’ errors.

### Timing and Experimental Conditions

We used a slow event-related design to prevent contamination of the blood oxygen level-dependent (BOLD) response by any potential artifacts generated by the hand movement. Trials were spaced every 18 s to allow the hemodynamic signal to return to baseline between trials. Each trial started with the Auditory Action-cue that consisted of a recorded voice that said “Grasp” or “Reach” and instructed the participant about the task to be performed at the end of the trial (**Figure 2A**). The auditory cue was followed by a delay of 8 s (Delay 1), after which the stimulus was illuminated for 250 ms showing the rod in one of the two orientations (Visual Orientation-cue). The visual cue consisted of the brief illumination of the 3D rod presented in one of two possible orientations: horizontal (approximately 0° angle) or vertical (approximately −45° angle). The visual cue was followed by another delay of 8 s (Delay 2), after which an auditory go cue instructed the participant to perform the action that had been instructed at the beginning of the trial. After 3 s, a “Beep” sound cued the participant to return the hand to the home position. One second later, the fixation light was turned off for 2 s during which participants were instructed to gaze freely. The same or different fixation light was then turned on and participants fixated the LED in the dark.

Note that the delays following the Action and Orientation-cue allowed us to examine the brain activation in the phase that followed action instruction and visual input while also allowing us to separate the sensory processing from the motor response.

Each run consisted of 8 trials and each experimental condition was presented one time in each run. The order of the conditions was randomized in each run. A baseline of 18 s was added at the beginning and at the end of each run yielding a run time of approximately 6 min per run. Each participant performed 6 runs, for a total of 6 trials per experimental condition. A session for one participant included set-up time (~45 min), six functional runs and one anatomical scan, and took approximately 90 minutes to be completed.

### Imaging Parameters

All imaging was performed at York University (Toronto, ON, Canada) using a 3-T whole-body MRI system (Siemens Magnetom TIM Trio, Erlangen, Germany). The posterior half of a 12-channel receive-only head coil (6 channels) at the back of the head was used in conjunction with a 4-channel flex coil over the anterior part of the head (see Monaco et al., 2014). The anterior part of the 12-channel coil was removed to allow the participant to see the stimuli directly and comfortably but at a cost of anterior signal loss, hence the addition of the 4-channel flex coil. The posterior half of the 12-channel coil was tilted at an angle of 20° to allow the direct viewing of the stimuli. We used an optimized T2-weighted single-shot gradient echo echo-planar imaging (211-mm field of view [FOV] with 64 × 64 matrix size, yielding a resolution of 3.3-mm isovoxel; 3.3-mm slice thickness with no gap; repetition time [TR] = 2 s; echo time [TE] = 30 ms; flip angle [FA] = 90°). Each volume comprised 38 slices angled at approximately 30° from axial (i.e., approximately parallel to the calcarine sulcus) to sample occipital, parietal, posterior temporal, and posterior/superior frontal cortices. The slices were collected in ascending and interleaved order. During each experimental session, a T1-weighted anatomical reference volume was acquired along the repeated orientation as the functional images using a 3D acquisition sequence (256 × 240 × 192 FOV with the repeated matrix size yielding a resolution of 1-mm isovoxel, inversion time, TI = 900 ms, TR = 2300 ms, TE = 5.23 ms, FA = 9°). The coil configuration used allowed coverage of most part of the brain, except for the ventral part of the cerebellum.

### Preprocessing

Data were analyzed using the Brain Voyager QX software (Brain Innovation 2.8, Maastricht, The Netherlands). Functional data were superimposed on anatomical brain images, aligned on the anterior commissure–posterior commissure line, and transformed into Talairach space (Talairach and Tournoux 1988). The first 2 volumes of each fMRI scan were discarded to allow for T1 equilibration. Functional data were preprocessed with spatial smoothing (full-width at half-maximum = 8 mm) and temporal smoothing to remove frequencies below 2 cycles per run. Slice-time correction with a cubic spline interpolation algorithm was also performed. Functional data from each run were screened for motion or magnet artifacts with cine-loop animation to detect eventual abrupt movements of the head. In addition, we ensured that no obvious motion artifacts (e.g., rims of activation) were present in the activation maps from individual participants. Each functional run was motion corrected using a trilinear/sinc interpolation algorithm, such that each volume was aligned to the volume of the functional scan closest to the anatomical scan. The motion correction parameters of each run were also checked: runs that showed abrupt head motion over 1 mm were discarded from further analyses.

### General linear model

To investigate which brain areas are involved in our task, we conducted voxelwise analyses on group data. For each participant, we used a general linear model (GLM) that included a predictor for each event of the trial. Each predictor was derived from a rectangular wave function convolved with a standard hemodynamic response function (HRF; Brain Voyager QX’s default double-gamma HRF). In particular, we used predictors covering the following time windows: 2 s (or 1 volume) for the Action-cue and Orientation-cue, 6 s (or 3 volumes) for the Delay 1 and Delay 2, 4 s (or 2 volumes) for the action execution, 4 s (or 2 volumes) for the free gazing and saccade. Our GLM included two predictors for the Action-cue and Delay 1 (Align and Grasp), four predictors for the Orientation-cue [(Vertical, Horizontal) x (Left of fixation, Right of fixation)], eight predictors for Delay 2 and Action execution [(Align, Grasp) x (Vertical, Horizontal) x (Left of fixation, Right of fixation)], one predictor for the Saccade.

The group random effect (RFX) GLM included 25 predictors for each participant. Contrasts were performed on the %-transformed beta weights (β).

### Voxelwise and ROI Analyses

We ran two voxelwise contrasts to test our hypotheses. Specifically, our first hypothesis was aimed to test if action intention modulates sensory processing (Hypothesis 1 in **Figure 2C**, left panel). To this aim, we examined whether the activation during the delay following the Visual Orientation-cue (Delay 2) would be influenced by the action that was instructed at the beginning of the trial. This would be reflected in higher activation for one action over the other during Delay 2. To test this hypothesis, we performed Contrast 1: Delay 2 (Align > Grasp). It is important to emphasize that the participants: 1) had received the auditory instruction about the action 10 s earlier, 2) were not performing any action yet and 3) were fixating the fixation LED in the dark during Delay 2. Surprisingly, this contrast elicited activation in the EVC despite the fact that participants were in the dark. In order to determine whether our area corresponds to V1, V2, V3, etc. we used a published probabilistic atlas (Wang et al. 2015) that provides a dataset with the full probability maps of topographically organized regions in the human visual system (www.princeton.edu/∼napl/vtpm.htm). In particular, the atlas provides the probabilistic maps generated from a large population of individual subjects (N = 53) tested with standard retinotopic mapping procedures and allows defining the likelihood of a given coordinate being associated with a given functional region for results obtained from any independent dataset once transformed into the same standard space. Therefore, we converted our Talairach coordinates in MNI space and used the atlas to examine whether the coordinates fall within V1, V2, V3, etc. The area in the EVC corresponds to the primary visual cortex (V1) in both hemispheres (**Supplementary Figure 1**).

Further, we ran an ROI analysis to localize the portion of the EVC that corresponds to the retinotopic location of the stimulus relative to the fixation point. This allows examining the effects in the localized region without biases related to non-independent selection criteria of the voxelwise analysis. To this aim, we ran the ROI contrast: Orientation-cue > Baseline and extracted the β weights from Delay 1, Delay 2 and Action execution from the localized ROI in the EVC. To examine whether the areas shown with Contrast 1 during Delay 2 were also shown during the execution phase we ran Contrast 2 to identify areas that showed higher activation for Align vs. Grasp during Action execution: Action execution (Align > Grasp). This contrast allowed us to identify areas whose activation reflects the bio-mechanical aspects of performing one action vs. the other.

Our second hypothesis was aimed at testing if the effects of the preparatory-set on sensory processing are orientation-specific (Hypothesis 2 in **Figure 2C,** right panel). To this aim, we performed Contrast 3: Delay 2 (Align Vertical > Align Horizontal).

Before testing our hypotheses, we explored the network of areas showing a general response in Delay 1 following the Auditory Action-cue, in Delay 2 following the Visual Orientation-cue, and during Action execution. We ran three contrasts on the group data and shown in **Figure 3**: A) [Delay 1 (Align + Grasp) > Baseline], B) [Delay 2 (Align + Grasp) > Baseline], and C) [Action execution (Align + Grasp) > Baseline]. As expected, a wide network of areas ranging from occipital to parietal and frontal cortex were involved in all three phases of the task. Strikingly, the EVC (including the posterior part of the Calcarine sulcus and the Cuneus) was also involved in all three phases despite the fact that there was no visual input at any of the stages analyzed here (Delay 1, Delay 2 and Action execution).

**Figure 3.**
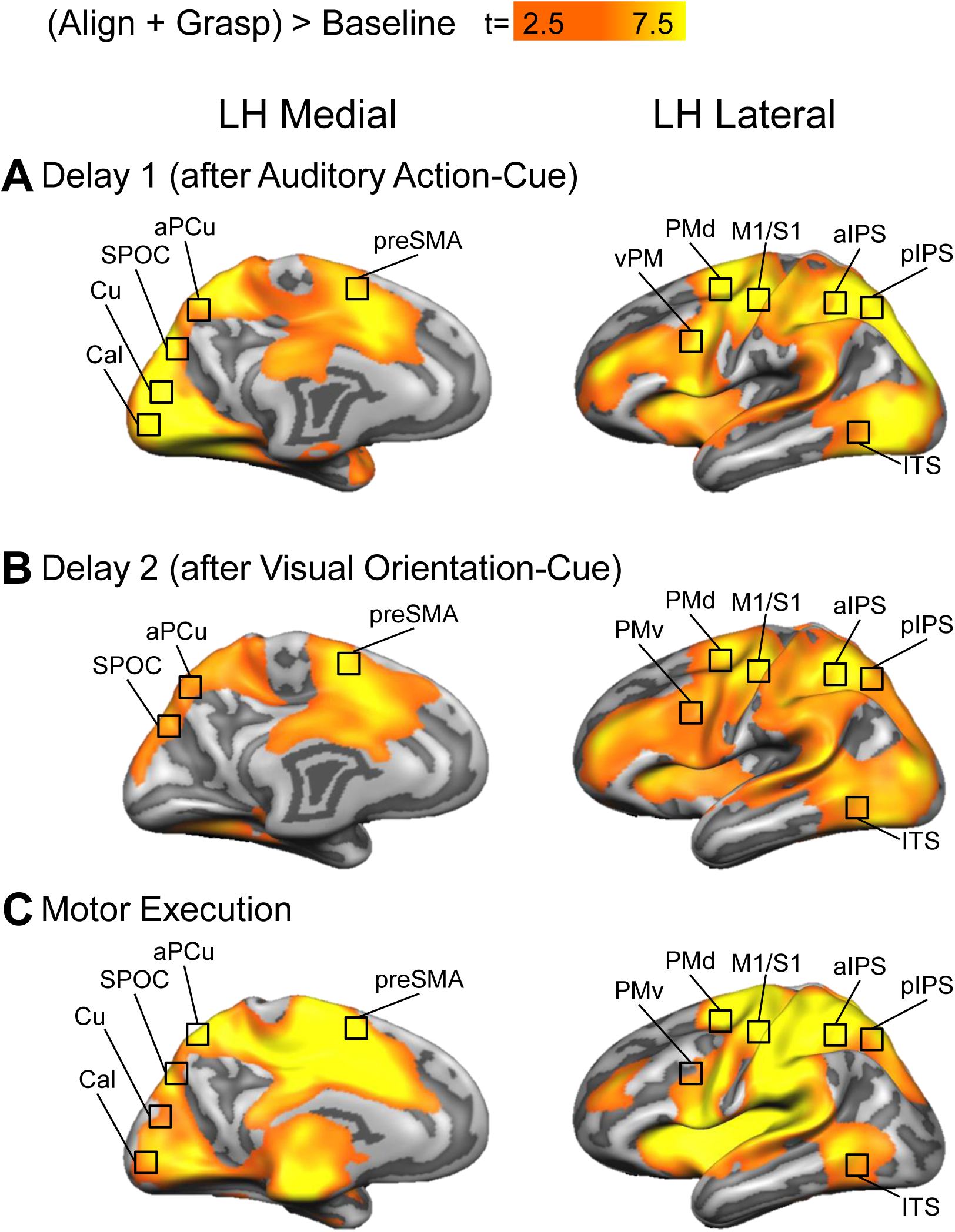
General activity for Align and Grasp conditions during the three phases of the trial in the medial and lateral view of the left hemisphere. Statistical parametric maps are overlaid on the average cortical surface derived from the cortex-based alignment performed on 11 of the 14 participants. The surface from the average of 11 subjects has the advantage to show less bias in the anatomy of the major sulci as compared to the surface of one participant only. We verified that activation from volumetric maps appeared on the corresponding region of the cortical surface. The statistical parametric maps shows areas with above-baseline activation for Align and Grasp conditions during: **A.** Delay 1 following the Auditory Action-cue (K= 69 for p<0.05)), **B. D**elay 2 following the Visual Orientation-cue (K= 52 for p<0.05), and **C.** the Action execution (K= 32 for p<0.05).

For each activation map generated by each contrast, we performed the cluster threshold correction (Forman et al. 1995) using the Brain Voyager’s cluster-level statistical threshold estimator plug-in (Goebel et al. 2006). This algorithm uses Monte Carlo simulations (1000 iterations) to estimate the probability of a number of contiguous voxels being active purely due to chance while taking into consideration the average smoothness of the statistical map. Because map smoothness varies with the contrast, different contrasts have different cluster thresholds. In cases in which the activation foci did not survive cluster threshold correction at an alpha-correction level of 0.05, we indicated the regions with a star. For each area, we extracted the β weights in each condition for further analysis. The minimum cluster sizes were estimated at 27 voxels of (3 mm)^3^ for a total volume of 729 mm^3^ for Contrast 1, 37 voxels of (3 mm)^3^ for a total volume of 999 mm^3^ for Contrast 2, 16 voxels of (3 mm)^3^ for a total volume of 432 mm^3^ for Contrast 3. The minimum cluster sizes for the contrasts A, B and C were estimated at 69, 52 and 32 voxels of (3 mm)^3^, respectively, for a total volume of 1863, 1404 and 864 mm^3^.

For areas defined by the voxelwise contrasts 1, 2 and 3, where the data were not selected by independent contrasts, the data graphs are based on non-independent contrasts. We have included these to show unambiguously the effects for the preparatory-set influences on sensory processing to enable the reader to easily see the pattern of results, including effects that are independent of the selection criteria. Contrasts that are non-independent are clearly indicated in square brackets.

### Statistical Analyses

For each activation map aimed to test our hypotheses we ran comparisons on the β weights from each area.

The first group of comparisons was aimed to test the nature of the modulation observed during Delay 2(Hypothesis 1, **Figure 2C**, left panel). In particular, we tested whether the observed effects truly reflected the action-driven modulation of sensory processing rather than a difference between instructed actions regardless of sensory processing. This would be shown by higher activation for Action 1 vs. Action 2 in Delay 2 but not Delay 1 or Action execution. To test this hypothesis, we ran three two-tailed paired-sample t-tests with a Bonferroni correction for three comparisons (P<0.0166). In addition, we tested whether the observed action-driven modulation was specific to Delay 2. This would be reflected in a larger difference in activation between Action 1 and Action 2 in Delay 2 vs. Delay 1, and Delay 2 vs. Action execution (Hypothesis 1, **Figure 2D**, left panel). For this purpose, we ran two two-tailed paired-sample t-tests with a Bonferroni correction for two comparisons (P<0. 025).

The second group of comparisons was aimed to test whether orientation-specific effects of action intention on sensory processing are specific to one action but not the other. Therefore, we performed two comparisons aimed to show whether the activation for Orientation 1 was higher than Orientation 2 in Action 1 but not Action 2 (Hypothesis 2, **Figure 2C**, right panel). To test this hypothesis, we ran two two-tailed paired-sample t-tests with a Bonferroni correction for two comparisons (P<0.025). Orientation-specific effects would also be reflected in a larger difference in activation between Orientation 1 and Orientation 2 in Delay 2 for Action 1 but not Action 2 (Hypothesis 1, **Figure 2D**, right panel). We tested this we ran two two-tailed paired-sample t-tests with a Bonferroni correction for two comparisons (P<0. 025).

To test possible effects of the visual hemifiled in which the stimulus was presented (Left and Right location relative to the gaze) in all areas we compared the activation for Left vs. Right stimulus presentation during Delay 2 and Action execution. This comparison did not yield any significant effect of visual hemifiled. Therefore we collapsed the data for Left and Right stimulus location relative to gaze.

Statistical differences are indicated on bar graphs (**Figures 5, 6 and 7**). To further illustrate the differences graphically with appropriate error bars for each effect, for each area, we computed differences in activation between conditions in Delay 1, Delay 2 and Action execution along with the 98.34% or 97.5% con?dence limits. We described statistical effects that are signi?cant at a corrected P-value, unless specified.

### Psychophysiological Interaction analysis (PPI)

We used the psychophysiological interaction method (Friston et al. 1997; McLaren et al. 2012; O’Reilly et al. 2012) to estimate the task-specific changes in effective connectivity between our seed region (LH V1) and the rest of the cortex. The PPI identifies brain regions whose functional connectivity is task-dependent and results from an interaction between the psychological component (the task) and the physiological component (the time course) of the seed region. In particular, we examined which brain areas show task-specific correlations with V1. With this aim, we created a PPI model for each run for each participant. The PPI model included: 1) the physiological component corresponding to the z-normalized timecourse extracted from the seed region, 2) the psychological component corresponding to the task model (boxcar predictors convolved with a standard hemodynamic response function), and 3) the psychophysiological interaction component, corresponding to the z-normalized timecourse multiplied, volume by volume, with the task model. The boxcar predictors of the psychological component were set to +1 for the Align task, −1 for the Grasp task, and zero for baseline. The psychological and physiological components were added as co-variates to our model to account for confounds related to 1) an effect of task, regardless of the physiological component, and 2) a correlation with the seed ROI timecourse, regardless of the task and shared task input. We selected our seed region based on the contrast of Align vs. Grasp in Delay 2. Therefore, areas that show a task preference similar to the seed region might show a positive relationship just based on the task and shared task input, regardless of whether these areas are functionally connected. In addition, a positive relationship with the seed region might be due to anatomical connections or shared subcortical inputs even in absence of task-specific effect. The motion correction parameters from each participant were added as co-variates of no interest. The individual GLM design matrix files were used for a random effects model analysis (Friston et al. 1999). The statistical threshold criterion was set to p < 0.05 for all presented contrasts, and the connectivity map was corrected for multiple comparisons suing the Monte Carlo simulation approach (Forman et al. 1995).

## Results

To test our hypotheses, we performed two voxelwise analyses aimed at testing the predictions illustrated in **Figure 2C and D**. Talairach coordinates and number of voxels for each area resulting from the activation map of the contrasts, are reported in **Table 1**. The activation maps and corresponding β weights for these voxelwise results are illustrated in **Figures 4 and 7**. Effects that are non-independent of the criterion used to select the voxels are clearly indicated in square brackets. Statistical values of the post hoc t-tests for each area are reported in **Tables 2 and 3**.

**Figure 4.**
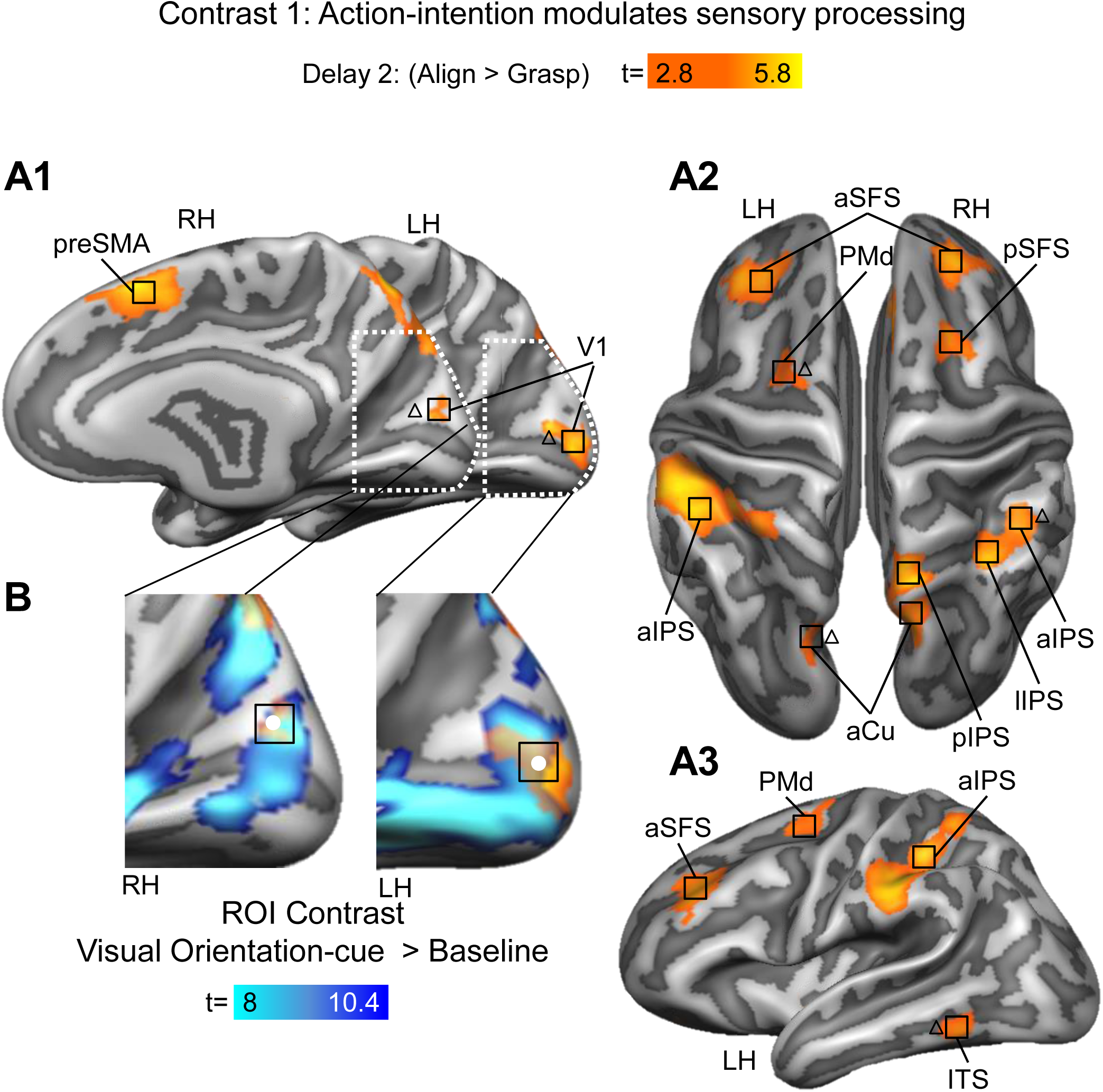
Influence of action intention on sensory processing. Statistical parametric maps are overlaid on the average cortical surface derived from the cortex-based alignment performed on 11 of the 14 participants. The surface from the average of 11 subjects has the advantage to show less bias in the anatomy of the major sulci as compared to the surface of one participant only. We verified that activation from volumetric maps appeared on the corresponding region of the cortical surface. **A1-3.** Statistical parametric map obtained with the RFX GLM showing the results for Hypothesis 1 (Contrast 1, K= 27 for p<0.016). **B.** The inserts show an enlarged view of the activation maps in the Calcarine sulcus in the left and right hemisphere. The activation for the visual presentation of the stimulus (ROI contrast: Visual Orientation-cue > Baseline) is shown in blue.

**Table 1.** Talairach coordinates and number of voxels

| Brain areas | Talairach coordinates |  |  | Number of voxels |
| --- | --- | --- | --- | --- |
|  | X | Y | Z |  |
| LH aSFS | -31 | 34 | 35 | 774 |
| RH aSFS | 24 | 39 | 35 | 627 |
| RH preSMA | 9 | 12 | 50 | 755 |
| RH pSFS | 20 | 5 | 53 | 505 |
| LH PMd | -25 | -8 | 57 | 501 |
| RH PMd | 24 | -4 | 48 | 507 |
| LH PMv | -35 | -6 | 36 | 101 |
| LH M1 | -41 | -21 | 50 | 991 |
| LH S1 | -44 | -30 | 54 | 871 |
| RH S1 | 42 | -29 | 53 | 697 |
| LH aIPS | -43 | -38 | 41 | 999 |
| RH aIPS | 42 | -43 | 40 | 567 |
| LH mPCS | -25 | -42 | 56 | 789 |
| RH mPCS | 18 | -43 | 56 | 419 |
| LH mIPS | -18 | -52 | 51 | 767 |
| LH IIPS | -23 | -46 | 39 | 232 |
| RH IIPS | 36 | -51 | 42 | 550 |
| RH pIPS | 12 | -62 | 55 | 774 |
| LH SPOC | -18 | -75 | 26 | 525 |
| RH SPOC | 18 | -74 | 27 | 598 |
| LH ITS | -53 | -53 | -11 | 475 |
| LH aPCu | -14 | -78 | 37 | 296 |
| RH aPCu | 11 | -62 | 41 | 517 |
| LH V1 | -3 | -89 | 1 | 228 |
| RH V1 | 1 | -75 | 9 | 374 |

Note: Area abbreviations as in figure legends.

**Table 2.**
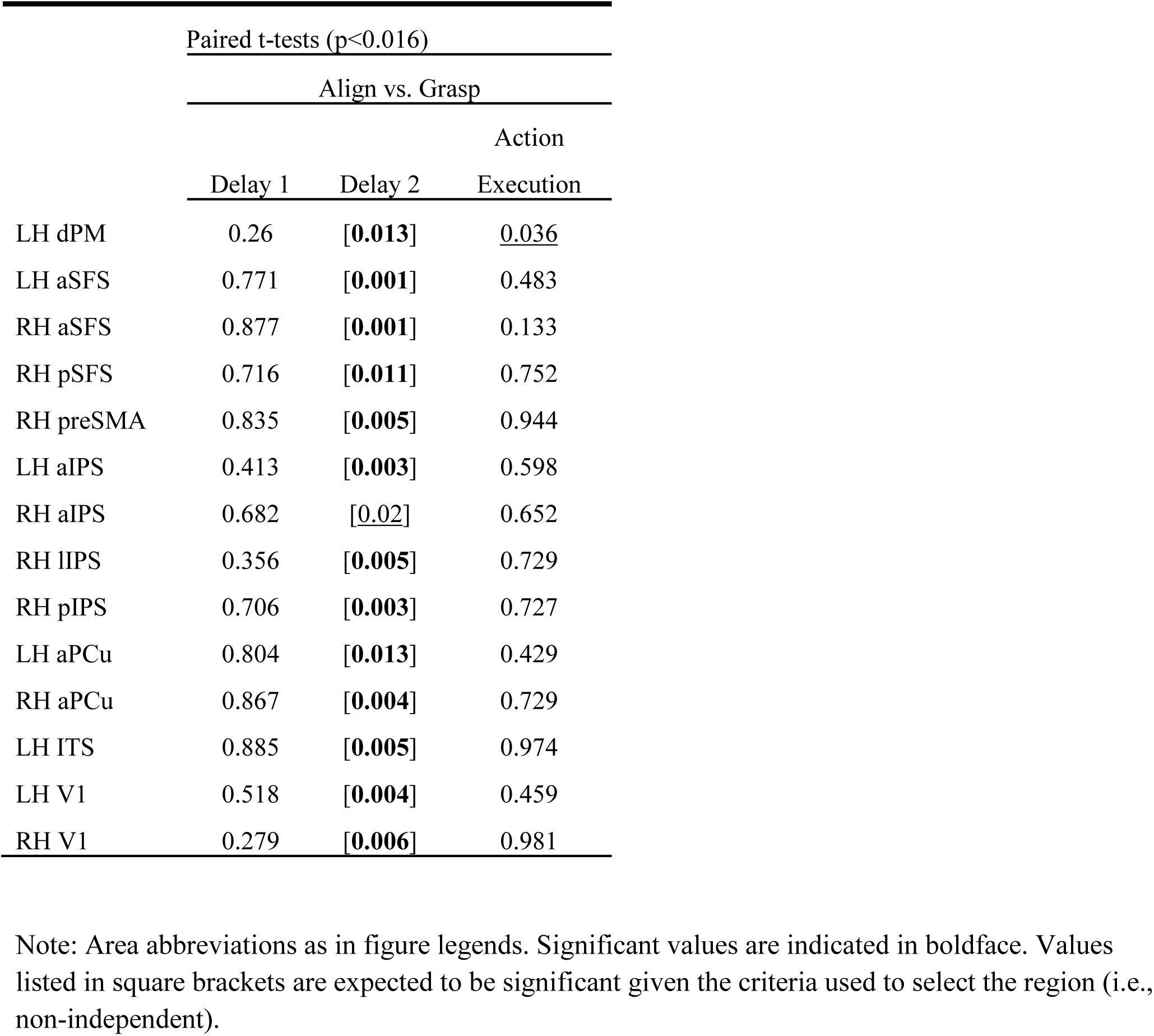
Statistical values for contrast 1

**Table 3.** Statistical values for contrast 2

| | Paired t-tests ($p < 0.016$) | | |
| --- | --- | --- | --- |
|  | Delay 1 | Delay 2 | Action Execution |
| LH dPM | 0.26 | **0.013** | [0.036] |
| RH dPM | 0.136 | 0.136 | [0.040] |
| LH M1 | 0.733 | 0.417 | [**0.001**] |
| LH S1 | 0.886 | 0.405 | [**0.001**] |
| RH S1 | 0.541 | 0.690 | [**0.002**] |
| LH mPCS | 0.870 | 0.142 | [**0.002**] |
| RH mPCS | 0.162 | 0.703 | [**0.007**] |
| LH mIPS | 0.173 | 0.338 | [**0.001**] |
| LH SPOC | 0.331 | 0.456 | [**0.012**] |
| RH SPOC | 0.724 | 0.415 | [**0.002**] |

Note: Area abbreviations as in figure legends. Significant values are indicated in boldface. Values listed in square brackets are expected to be significant given the criteria used to select the region (i.e., non-independent).

In addition, we tested whether the visual field in which the stimulus was presented affected the modulation of sensory information induced by action intention during Delay 2, when the stimulus was no longer visible. Given the lack of any significant visual field effect in our areas of interest, we collapsed the data for Left and Right stimulus location relative to the gaze. However, the Left and Right presentation of the stimulus during the Visual orientation-cue did elicit the expected contralateral activation in the visual cortex (not shown).

### General Observations: Preparatory Set, Planning, and Motor Execution

As an overview, our tasks (Align and Grasp) elicited above baseline activation in cortical areas involved in visual processing for action during Delay 1, Delay 2 and Motor execution (**Figure 3**; See **Supplementary Figure 1** for right hemisphere). These included well known reach and grasp areas such as the superior parietal occipital cortex (SPOC), anterior intraparietal cortex (aIPS), dorsal premotor cortex (PMd) and the pre-supplementary motor area (preSMA). It is noteworthy that non-motor areas, such as occipital areas involved in early vision (such as the Calcarine sulcus; Cal) and temporal areas (such as inferior temporal sulcus; ITS) known to be involved in higher level functions, were also activated during Delay 1 and Motor execution epochs despite the absence of visual stimulation.

Further, at this level of analysis many of these cortical areas were active throughout the duration of the task, including the first delay after auditory instruction (**Figure 3A**), the second delay after presentation of the visual stimulus (**Figure 3B**), and the final motor response (**Figure 3C**). This suggests that as soon as participants were instructed about the nature of the forthcoming action (Align or Grasp), preparatory activity (sometimes called ‘preparatory set’) commenced in these cortical areas. According to our experimental design, this preparatory activity would then combine with visual information about object orientation to allow the creation of a specific motor plan in Delay 2, which precedes motor execution. This is the focus of the remainder of our analysis, i.e., how specific action instructions and preparatory activity from Auditory Action-Cue and Delay 1 influence cortical visual processing for the purpose of motor planning in Delay 2.

### Testing Hypothesis 1: Action-specific enhancements of orientation processing

As noted above, our first hypothesis predicts that areas that differentially process visual information as a function of previous task instruction would show more activation for the Align than the Grasp task in Delay 2 (**Figure 2 C and D,** left panels). This effect would be shown with Contrast 1: (Align > Grasp) during Delay 2. **Figure 4 A1-3** provides the activation maps corresponding to this contrast. Note that during Delay 2 the stimulus was no longer visible. Contrast 1 yielded activation in bilateral primary visual cortex (V1, see Methods for localization of V1), anterior superior frontal sulcus (aSFS), anterior Cuneus (aCu), right posterior superior frontal sulcus (pSFS), pre-supplementary motor area (preSMA), lateral intraparietal sulcus (lISP), and posterior intraparietal sulcus (pIPS) as well as PMd and ITS. Thus, as predicted by Hypothesis 1, we found that a broad constellation of areas extending from occipital to temporal, parietal, and frontal cortex was modulated by the previous action instruction in Delay 2.

Higher activation in V1 for Align than Grasp conditions during Delay 2 was particularly surprising given the fact that: 1) participants were in the dark, 2) participants were not performing any movement and 3) the orientations of the stimuli were equally balanced for Align and Grasp conditions. Therefore, we analyzed this region in more depth. To confirm that the area identified by our contrast is V1, we used a probabilistic atlas (Wang et al. 2015) that shows that our regions falls within the boundaries of V1 in both hemispheres (**Supplementary Figure 1**, see Methods for details). To test the effects in V1 in an independent manner and avoid circularity issues, we localized the retinotopic location of the oriented rod in the EVC using the ROI contrast: Visual Orientation-cue > Baseline. The activation map in **Figure 4B** shows the occipital areas that were active during the visual presentation of the stimulus in the occipital cortex. The activation above the Calcarine sulcus is consistent with the location of the stimulus below the fixation point. This area overlaps with the area identified with Contrast 1 and statistical analyses on the β weights extracted from either area yielded similar results. Thus, surprisingly, even V1 was active and modulated by action instruction during Delay 2, even if participants were lying still in the dark.

One explanation for these results is that the auditory action instruction produced task-specific modulations of cortical activity that lasted through Delay 1 and Delay 2, independent of visual input. Therefore, to completely fulfil our hypothesis these areas would have to show task-dependent modulations that are specific to Delay 2. **Figure 5** provides the difference in β weights as shown in **Figure 2D** (left panel). For the unsubtracted β weights, see **Supplementary Figure 3**. As expected for this contrast, **Figure 5** indicates that all areas showed higher activation for Align than Grasp conditions in Delay 2, (p-uncorrected in aIPS in the right hemisphere). Importantly, the difference in activation between Align and Grasp conditions was higher in Delay 2 than Delay 1 for all areas except for the aSFS and PMd in the left hemisphere. This difference was significant at p-uncorrected in left V1, as well as right aSFS, aIPS and bilateral aCu. These results suggest that the effect of the previous action instruction in Delay 2 reflects a specific influence in processing the orientation of the visual stimulus for motor planning, rather than a non-specific difference related to the action instruction. In addition, there was also higher difference between Align and Grasp conditions in Delay 2 than Action execution in V1, left aIPS, right aCu and aSFS.

**Figure 5.**
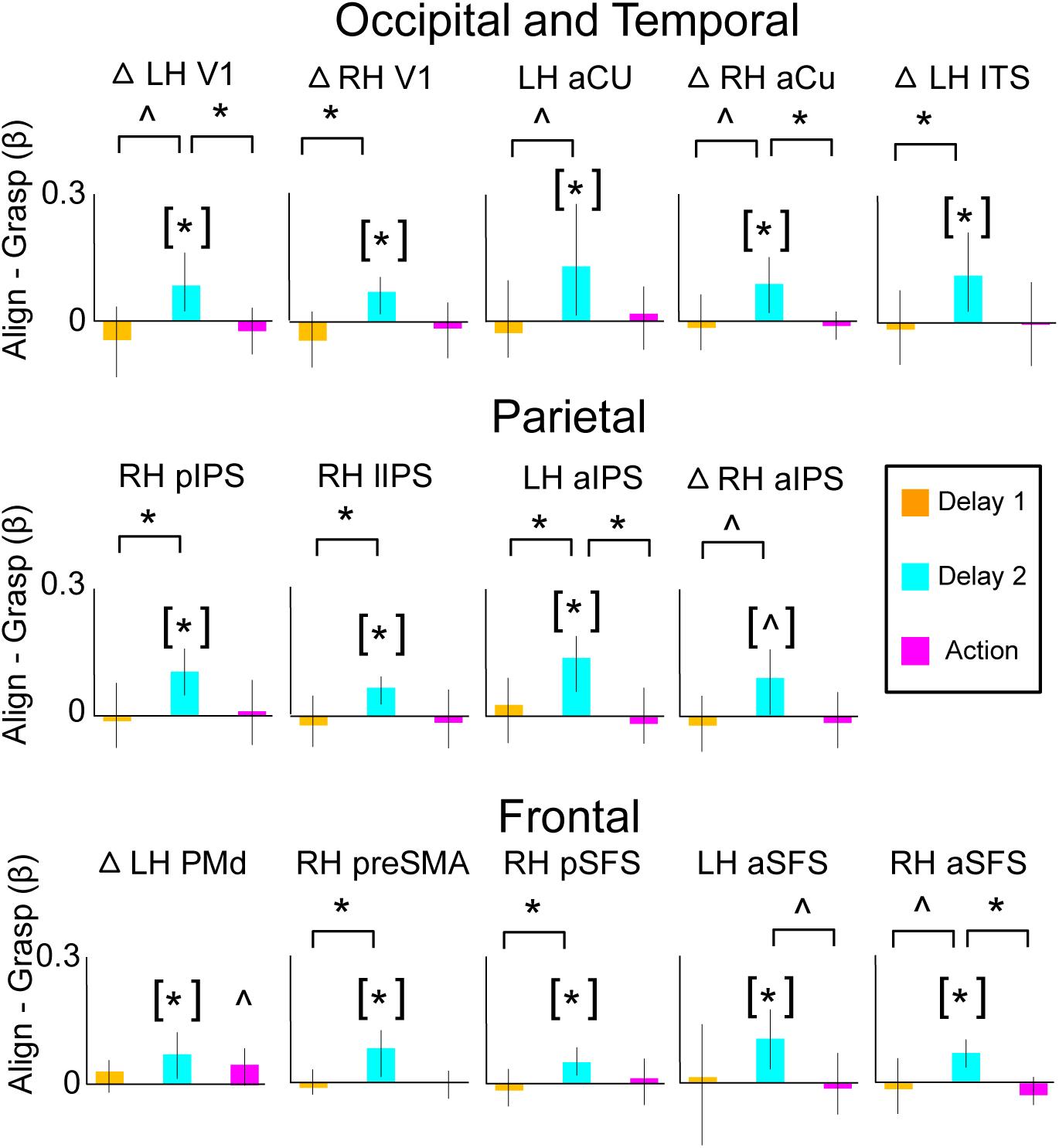
For each area identified with Contrast 1 and shown in A1-3, bar graphs indicate the difference in β weights for the two tasks (Align and Grasp) during the three phases of the task (Delay 1, Delay 2 and Action execution). *Significant statistical differences among conditions (p corrected). ^ Significant statistical differences among conditions (p uncorrected). Square brackets indicate significant effects that are non-independent given the criteria used to select the region.

Aligning the hand vs. grasping an object must require different motor commands, so it would be surprising if all task-specific activation ceased during the movement (**Figure 5** shows no difference between Grasp vs. Align during the Action). To test if other, different brain areas show task-specific motor activation during the action, and thus provide a ‘double dissociation’ for our hypothesis, we performed Contrast 2: (Align > Grasp) during Action execution. **Figure 6A** shows the activation map for this contrast, yielding a constellation of areas including SPOC, medial intraparietal sulcus (mIPS), medial post-central sulcus (mPCS), primary somatosensory cortex (S1), primary motor cortex (M1), and PMd, known to be involved in manual motor control. **Figure 6B** shows the corresponding differences in β weights for these areas, arranged across Delay 1, Delay 2, and Action execution. One can see that (with the exception of PMd) most of these areas are different from the areas found with Contrast 1 (**Figure 4**) and instead show task-related activation that is specific to the motor execution phase. Thus, we found a double dissociation showing a set of areas involved in task-specific planning activity in response to the visual stimulus during Delay 2 (**Figures 4**) vs. a set of areas that show task-specific motor activity during the movement (**Figure 6**). This shows that, although overall activation appeared to be fairly consistent across our delays and action execution (**Figure 3**), the details of cortical processing were different in each phase.

**Figure 6.**
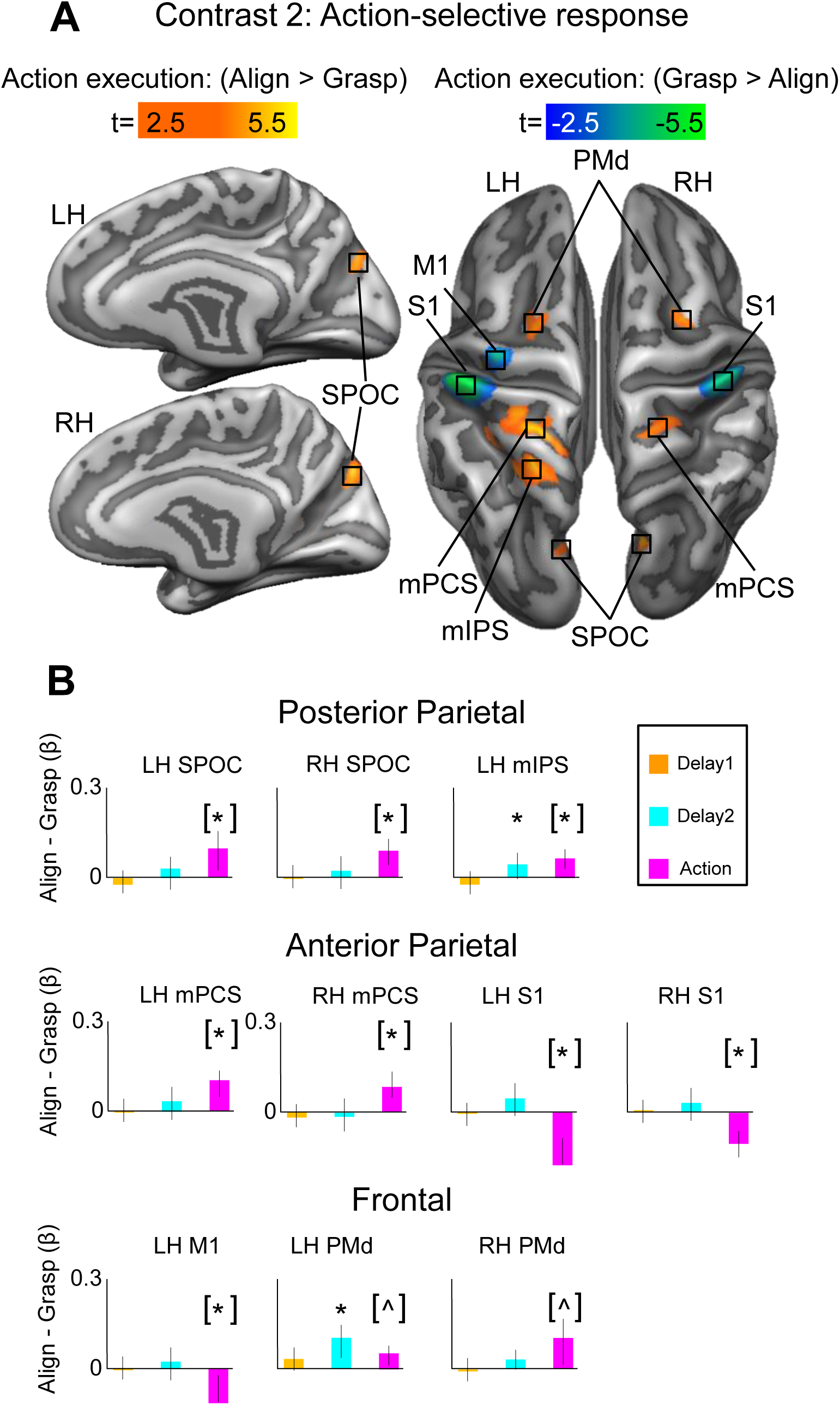
Action execution. **A.** Statistical parametric map obtained with the RFX GLM showing areas with higher activation for Align than Grasp tasks during the Action execution phase (Contrast 2, K= 37 for p<0.016). **B.** Bar graphs indicate the β weights for the two tasks (Align and Grasp) during the three phases: Delay 1, Delay 2, Action execution. *Significant statistical differences among conditions (p corrected). ^ Significant statistical differences among conditions (p uncorrected). Square brackets indicate significant effects that are non-independent given the criteria used to select the region.

### Testing Hypothesis 2: Task-specific enhancements in visual orientation selectivity

Recall that Hypothesis 2 predicts that the Align vs. Grasp instruction will have differential effects on object orientation specificity during Delay 2 (**Figure 2 C and D,** right panels), with the expectation that the Align task would require higher orientation-specific processing than the Grasp task, as the whole hand had to be adjusted according to the orientation of the rod. If this is the case, one orientation would elicit higher activation than the other for the Align but not the Grasp condition during Delay 2. To test this hypothesis, we ran Contrast 3: (Align Vertical > Align Horizontal) during Delay 2. As shown in **Figure 7A**, this contrast yielded higher differential orientation responses in four areas known to be involved in planning manual action: bilateral lateral intraparietal sulcus (lIPS), left ventral premotor cortex (PMv), and right PMd. The corresponding contrast for the Grasp condition: (Grasp Vertical > Grasp Horizontal) during Delay 2, yielded no significant results. Importantly, the difference between Vertical and Horizontal orientation during Delay 2 is higher in Align than Grasp conditions (**Figure 7B**). Thus, these three areas conformed to our hypothesis of task-specific visual orientation specificity during motor planning (**Figure 2D,** right panel)

**Figure 7.**
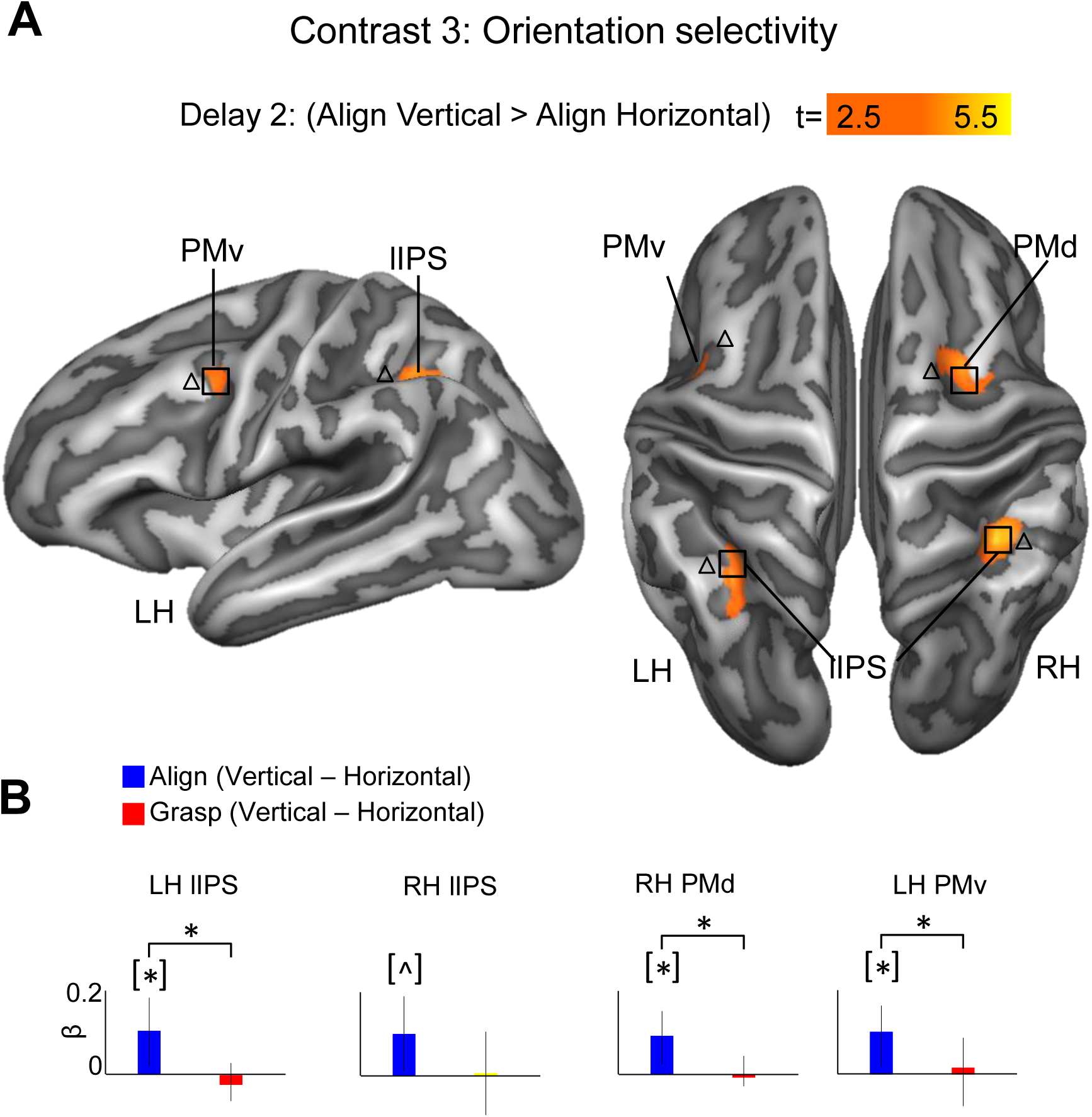
Orientation selectivity during Delay 2. **A**. Statistical parametric map obtained with the RFX GLM showing the effect for Hypothesis 2 (Contrast 3, K= 16 for p<0.025). **B.** Bar graphs indicate difference between the β weights for the two tasks (Align and Grasp) in the two orientations (Vertical and Horizontal) during Delay 2. *Significant statistical differences among conditions (p corrected). ^ Significant statistical differences among conditions (p uncorrected). Square brackets indicate significant effects that are non-independent given the criteria used to select the region.

### Psychophysiological Interaction analysis

The results described above, in particular the action-specific modulation in V1, suggest that action-specific visual enhancements are mediated by reentrant pathways from motor areas. To test this hypothesis, we performed Psychophysiological Interaction analysis focusing on the connectivity of V1 as the seed region. We chose V1 because it should carry the earliest visual signals at the cortical level, and this would allow us to test the most extreme case of our hypothesis on reentrant pathways from higher-level areas to primary sensory areas. Specifically, we investigated which areas show enhanced connectivity with the left V1 during Delay 2 for Align as compared to Grasp conditions. The PPI results are illustrated in **Figure 8,** which shows bilateral preSMA, M1/S1 as well as the ITS and V1 in the left hemisphere. The M1/S1 region was found in the fundus of the Central Sulcus. Thus, V1 showed functional connectivity both with hand motor areas and the ventral visual stream during Delay 2. This analysis cannot show the directionality of influence, but note that during this delay there is no visual stimulus, so the enhanced connections between V1 and somatosensory and motor areas could not be driven by purely feedforward sensory signals from V1. Instead, this suggests the existence of a reentrant loop. Thus, this provides a potential neural substrate for reentrant influence of motor and higher visual signals on visual processing as early as V1.

**Figure 8.**
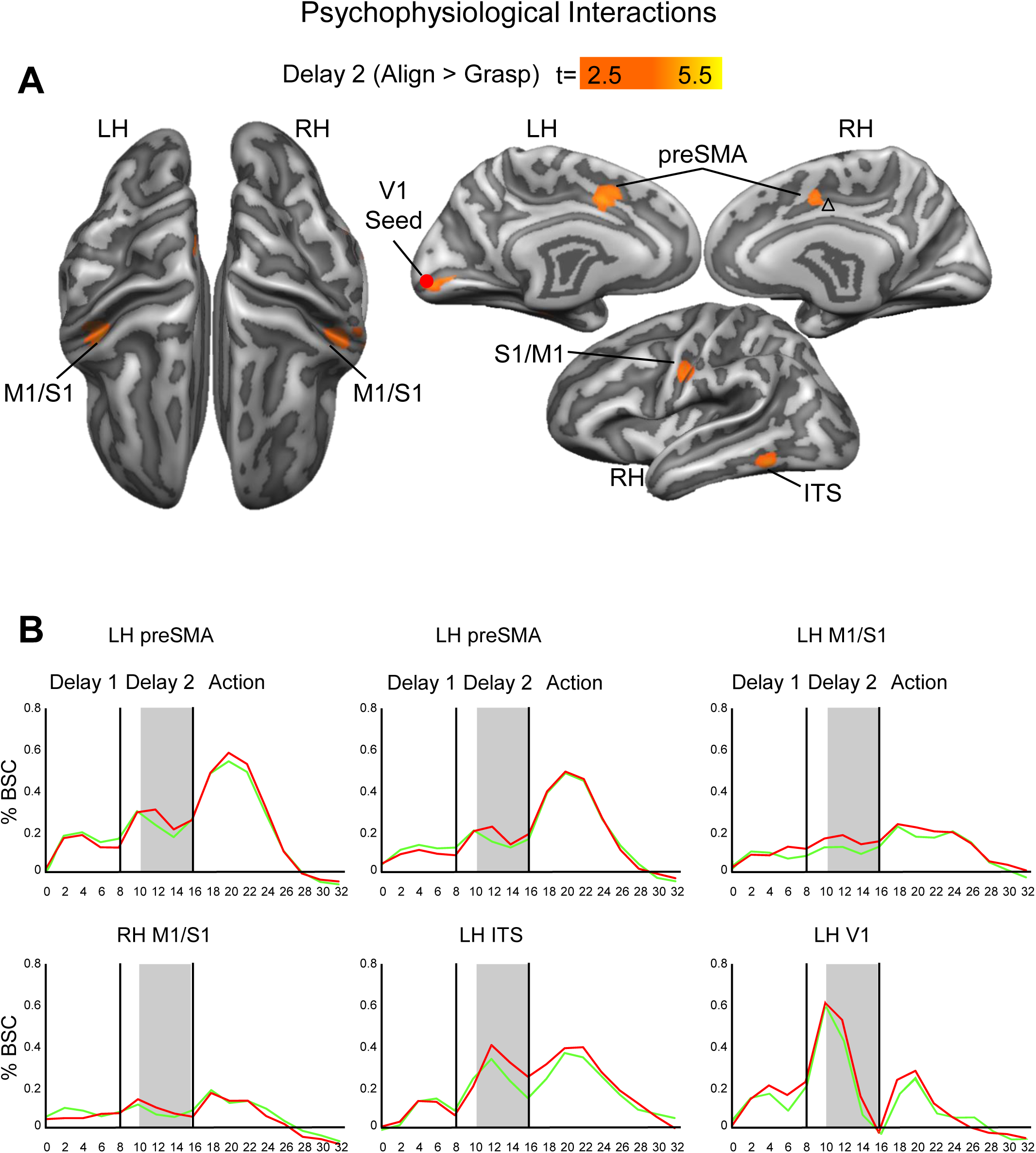
Functional Network of areas involved in the modulation of sensory processing driven by action intention. **A.** Statistical parametric map showing the psychophysiological interaction results using V1 in the left hemisphere as the seed region (K= 15 for p<0.05). **B.** Timecourse of the areas that functionally connected to V1 in the left hemisphere for processing Align vs. Grasp movement plans during Delay 2. The red circle corresponds to the peak voxel of the seed ROI.

## Discussion

Our results provide two novel findings. First, during the planning of manual action, the cortical processing of visual features depends on specific task instructions. As hypothesized, we observed instruction-specific (Align vs. Grasp) modulations in overall activation (**Figures 4 and 5**) and stimulus orientation specificity (**Figures 7**) in a constellation of cortical areas spanning V1, temporal, parietal and frontal cortices, including areas involved in hand actions (aIPS, SPOC, PMd). These patterns only emerged after the full set of information required to execute the movement became available, and were specific to the planning period after the presentation of the stimulus and before the “go” signal for action. Conversely, the go signal also evoked task-specific motor activation (**Figure 6**), but in a group of cortical areas most of which were not modulated during planning. Second, during the planning phase V1 showed stronger functional connectivity with motor areas (preSMA, M1/S1), as well as higher level visual cortex in the ventral stream (ITS) for the Align than Grasp condition. Indeed, the Align but not the Grasp task required participants to adjust the posture of the hand and wrist according to the orientation of the rod. Note that since these activations and task-specific functional connections occurred in the complete absence of light, well after visual stimulation and before the actual movement, our data suggest that they reflect reentrant feedback loops from frontal and parietal areas as well as higher order visual areas to the primary visual cortex.

Our results provide evidence that action intention modulates the response to sensory processing in the primary visual cortex and extend previous findings about the role of this area during execution. Indeed, previous investigations have reported re-activation of the EVC *during* the execution of actions in the dark towards unfamiliar shapes (Singhal et al., 2013) as well as the representation of certain object features for grasping, such as object elongation (Fabbri et al. 2016). Our results extend these findings by showing; 1) action-specific modulation in the primary visual cortex during action planning despite the lack of visual information, and 2) this processing is propagated and elaborated throughout the dorsal and ventral visual stream. Thus our results provide evidence that the role of the primary visual includes not only visual processing and memory, but also action-specific planning.

### Possible mechanisms for the influence of action instruction

We provide a framework for understanding the likely neural events that occurred during our experiment, embedding the current findings within a speculative model based on the literature (**Figure 9**). In order to organize this discussion, the task has been broken down into four phases, beginning with action instruction (**Figure 9A**) and its influence on preparatory set (**Figure 9B**). Clearly, the auditory instruction (Grasp and Align) initiated the preparatory activity that we observed during Delay 1 (**Figure 3A, Supplementary Figure 1**). This included widespread activation in well-known prefrontal and parietal-frontal areas primed for action, as well as extensive occipital-temporal activation, presumably primed for the expected visual stimulus. One can safely assume that the instruction was processed by auditory cortex (**Figure 9A**), which in some species has direct connections to primary visual cortex (Budinger et al. 2006; Campi et al. 2010). The instruction-dependent influence, most likely mediated by the language-processing areas (Broca’s area and Wernicke’s area) and executive control mechanisms, might have acted as a ‘trigger’. The premotor cortex is thought to play a role in ‘preparatory set’ (pre-cued activation of neurons in anticipation of action), possibly initiating the reentrant propagation of signals through motor-to-sensory areas. This likely included general priming of feature-specific processing mechanisms (Martinez-Trujillo and Treue 2004; Heuer, Schubö, et al. 2016), in this case intention-related orientation selectivity (Perry and Fallah 2017). However, we did not observe any action-specific responses during this first delay (**Figures 5 and 6**), or in the initial visual response before the second delay (not shown). It is likely that a reverberating circuit maintained the activation observed in Delay 1. Once set up, this circuit would readily program the manual response to the visual stimulus with a suitable action.

**Figure 9.**
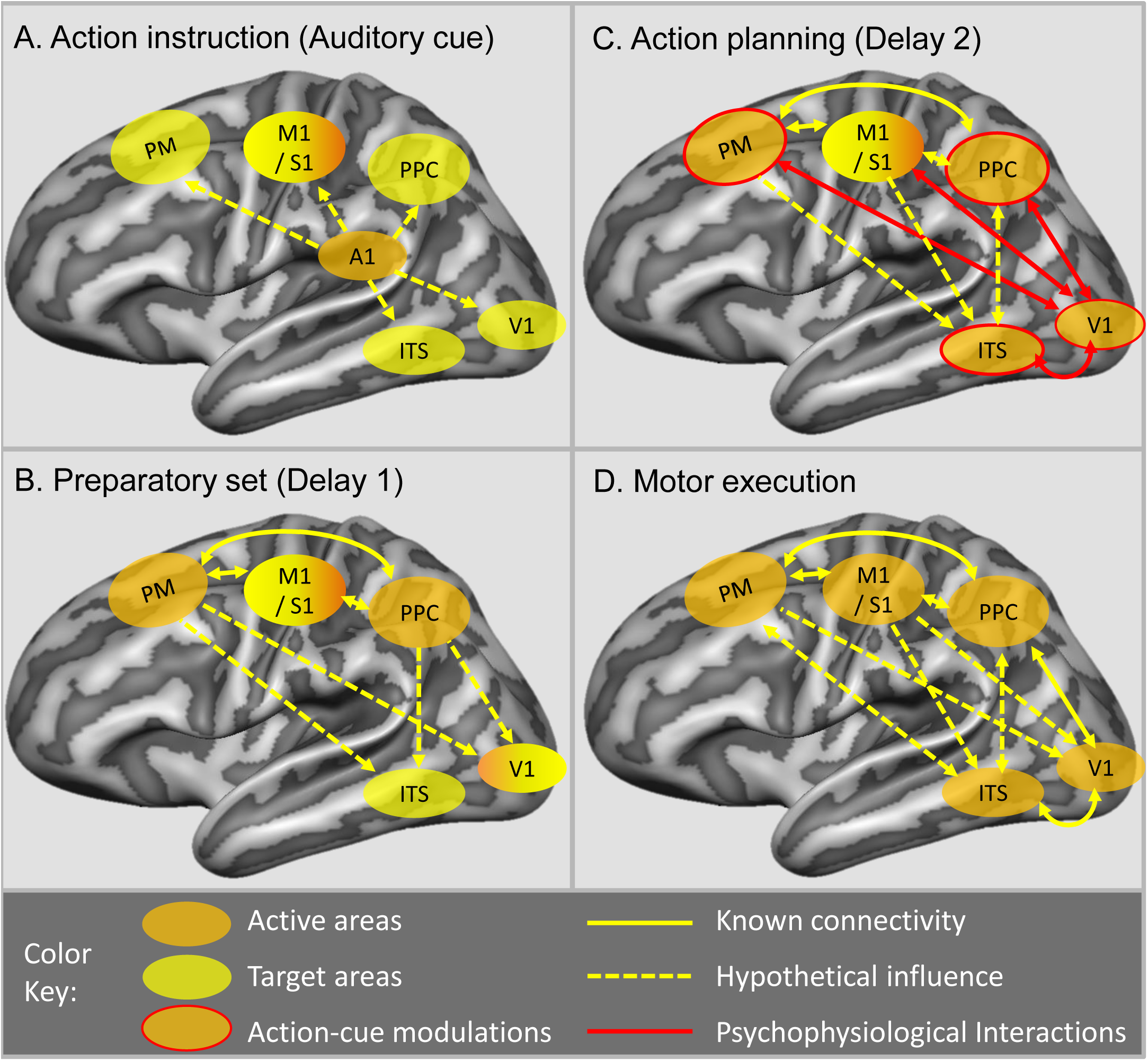
Schematic representation of functional networks involved in different stages of the experiment. While visual, occipito-temporal, parietal, somatosensory, motor and premotor cortices respond to action and orientation cues, reciprocal connections between all these areas are likely to occur during the planning phase preceding the movement to determine predictive coding activity in V1. In particular, we suggest that: **A.** the Auditory Action-cue primes motor preparation and upcoming object processing, **B.** Motor preparation primes the visual system for processing action-related object properties, **C.** Task-Dependent Integration of Visual Stimulus into motor plan, and **D.** Integrated network contributes to the execution of the movement.

### Instruction-Dependent Integration of the Visual Signal into the Motor Plan

Once the visual stimulus was shown it became possible for our participants to plan a specific movement with a specific orientation (**Figure 9 C**). This would involve the feedforward propagation of highly precise retinal input with the less precise and broader information conveyed to V1 through cortical feedback (Muckli et al. 2013; Chong et al. 2016; Petro and Muckli 2016). The second type of information participates to building and internal model that allows the brain to make predictions and anticipate the outcome of a movement generated by the self. Given the absence of online visual, somatosensory and motor information during the second delay, our results tackle the mechanisms underlying predictive coding and show that such process happens as early as in V1.

During Delay 2, we observed action and orientation selective responses in well-known visual areas (V1, ITS) and reach and grasp planning areas (pIPS, aIPS, PMd, preSMA). Since this specificity was not observed until the second delay period, it appears that neither instruction nor vision alone were sufficient to produces this pattern. Rather, it appears that preparatory set and visual stimulation were combined to produce action and orientation specific planning responses. Our psychophysiological interaction analysis suggests that the early modulations observed in primary cortex could be mediated by connections from preSMA and S1/M1, which is known to encode specific grasp before an actual movement (Gallivan, Mclean, Valyear, et al. 2011) and have interconnections with V1 (Miller and Vogt 1984).

Since action potentials have not been reported in V1 in the absence of vision, it is possible that at the top-down influence that we detected in occipital cortex is generated by subthreshold post-synaptic activity (Logothetis 2003), possibly intended for the attentional modulation of (normally available) visual input (Moore and Armstrong 2003). This interpretation would be consistent with evidence for BOLD activation, but not action potentials, in early visual areas during working memory tasks (Harrison and Tong 2009; Leavitt et al. 2017). However, as one proceeds to the higher levels of parietal and premotor cortex there is ample evidence for delay-related action potential activity (neurophysiology: (Snyder et al. 1997); fMRI: (Gallivan, Mclean, Smith, et al. 2011), so one would expect to see task-dependent modulations of action potentials in the areas we identified.

Importantly, area V1, aCu, aIPS and aSFS unequivocally show that the difference in activity between the Align and Grasp tasks is higher in the second delay than either the first delay or the Action execution phase. This shows that these instruction-dependent modulations were neither related to general action preparation or action execution, but rather were specific to processing visual input for the purpose of an action plan. Likewise, our finding of functional connectivity between visual and motor areas in the absence of either vision or movement suggests that this process is sub-served by reentrant connectivity. Therefore, it appears that action intention, likely initiated in the frontal cortex, shapes the way that vision is processed for action execution. However, when it comes time to act, our results show this this profile of action selectivity shifts to a different set of cortical areas, as described in the next section.

### Action execution

During Action Execution (Figure 9D) We found higher activation for Align than Grasp actions in areas known to have a crucial role in orienting the hand during actions, such as SPOC and PMd. Specifically, the superior parietal cortex of humans (SPOC), the putative homologue of macaques’ V6/V6A, is involved in orienting the hand during grasping movements (humans: Monaco et al., 2011, macaques: Fattori et al., 2009). In addition, SPOC has been shown to be crucial for the control of hand posture during grasping actions as lesions to this area in macaques lead to profound impairments in orientating the wrist and hand posture during grasping movements (Faugier-Grimaud et al. 1978; Battaglini et al. 2002). Similarly, area PMd in humans shows an involvement in hand posture (Monaco et al. 2011), and stimulation to this area disrupts adjustments in wrist orientation (Taubert et al. 2010).

We observed comparable activation for grasp and align movements in area aIPS during the execution of the action. This might seem at odds with the well-known role of area aIPS in grasping actions. One possible explanation is that the coarse whole hand grasp used in our experiment falls within the range of grasping movements that elicit the lowest activation in aIPS among the different types of grip. Indeed, a recent study shows that the activity in human aIPS is modulated by the amount of precision required by a movement. Specifically, grasping movements that require higher precision elicit higher activation than coarse grasping movements in the aIPS (Cavina-Pratesi et al. 2017). In addition, neurophysiological studies have shown the involvement of the aIPS in hand orientation (Murata et al. 2000; Baumann et al. 2009). These factors might have led to the lack of higher responses for grasp than align in area aIPS.

The primary somatosensory and motor area (S1 and M1) were the only areas that showed higher response for Grasp than Align during movement execution. This is likely due to the larger somatosensory feedback and finer motor output associated with grasping movements. In particular, two factors might have contributed to these results. First, during grasp but not align movements, the participants touched the object with the five digits eliciting a rich somatosensory stimulation. Second, there was a higher degree of online motor control of the five fingers in grasping but not aligning movements during action execution. Indeed, the position of the digits could be adjusted after movement initiation and once the fingers touched the object. In contrary, to properly align the hand to the orientation of the rod, the hand posture had to be defined prior to initiating the movement. Therefore, online movement adjustments were required to a lesser extent for Align than Grasp movement.

### Interactions with Perception

To this point we have focused on the dorsal visual stream of visuomotor control, but the ITS – a ventral stream area implicated in perception— also showed a prominent role in our analysis. This included general preparatory activity (**Figure 3**), action selectivity (**Figures 4 and 5**) and increased functional connectivity with V1 for Align than Grasp conditions during Delay 2 (**Figure 8**). Perception and action clearly interact: we act on objects that we perceive, and manual action produces various perceptual effects in vision. What then do our results say about this?

The ITS is associated with high-level object perception and interconnects with other areas of visual cortex. A recent neuroanatomy investigation on macaques has shown direct connections between associative parietal areas in the inferior parietal lobule and V1 and V2 as well as the TEO (Borra and Rockland 2011). Recent fMRI studies of functional connectivity in humans have also shown functional connections between areas in the dorsal stream and occipito-temporal cortex LOC (Hutchison and Gallivan 2016; Tal et al. 2016). Our results extend these findings and suggest that connectivity between V1 and a network of premotor, motor, and ventral stream areas interacts in order to provide task-specific perceptual information into the relevant motor plan.

The effective connectivity between V1 and motor areas observed here might also sub-serve visual and somatosensory-motor associations in subjective experience. For example, during manipulation we often visualize both the hand and object. This interpretation is in line with a recent study by Smith and Goodale (2015) showing that the activity pattern in S1 can be used to decode categories of objects that were only seen by the participant, with higher decoding accuracy for familiar objects.

In summary, our results suggest that the intent for specific actions could influence both conscious perception, and in turn influence how perception shapes action.

### Concluding remarks

The novel results of this study are that the intent for specific actions can influence visual processing as early as primary visual cortex (extending through temporal, parietal, and frontal cortex) and that this seems to originate from reciprocal functional connectivity between motor and sensory areas. This suggests that specific action plans shape how the brain both perceives and uses what the eyes see.

## Conflict Interest

The authors declare there are no conflicts of interest.

### Acknowledgements

We would like to thank Joy Williams for assistance during fMRI data collection, Saihong Sun, and Vittorio Iacovella.

## Funding

This work was supported by the Canadian Institute of Health Research and a Canada Research Chair award to J. D. Crawford (grant number 12736) and the European Union’s Horizon 2020 research and innovation programme under the Marie Sklodowska-Curie grant agreement No 703597 to Simona Monaco.

Author Contributions Statement: Simona Monaco designed the experiment, collected and analyzed the data, wrote the manuscript. Ying Chen collected the data. Doug Crawford: wrote the manuscript.

## Figure legends

*Supplementary Figure 1.* **Retinotopic visual areas.** The activation maps were taken from a published and freely available probabilistic atlas (Wang et al., 2014). The white circle shows the MNI coordinates of the occipital areas found with contrast 4 and corresponding to V1.

*Supplementary Figure 2*. Same as in Figure 3. Here we show the dorsal view of left and right hemisphere as well as the medial and lateral view of the right hemisphere.

*Supplementary Figure 3*. **Influence of action intention on sensory processing: Activation levels and statistical comparisons between the key conditions** for areas shown in Figure 4. for each area identified with Contrast 1 and shown in Figure 4, bar graphs indicate the β weights for the two tasks (Align and Grasp) during the three phases (Delay 1, Delay 2, Action execution).

*Supplementary Figure 4*. **Action selective responses: Activation levels and statistical comparisons between the key conditions** for areas shown in Figure 6. Figure legend as in Supplementary Figure 3.

*Supplementary Figure 5*. **Orientation selectivity: Activation levels and statistical comparisons between the key conditions** for areas shown in Figure 7. for each area identified with Contrast 3 and shown in Figure 7, bar graphs indicate the difference between the β weights for Vertical vs. Horizontal conditions in the two tasks (Align and Grasp) during Delay 2.

